# High-dimensional single-cell cartography reveals novel skeletal muscle resident cell populations

**DOI:** 10.1101/304683

**Authors:** Lorenzo Giordani, Gary J. He, Elisa Negroni, Hiroshi Sakai, Justin Y. C. Law, M. Mona Siu, Raymond Wan, Shahragim Tajbakhsh, Tom H. Cheung, Fabien Le Grand

## Abstract

Adult tissue repair and regeneration require the activation of resident stem/progenitor cells that can self-renew and generate differentiated progeny. The regenerative capacity of skeletal muscle relies on muscle satellite cells (MuSCs) and their interplay with different cell types within the niche. Yet, our understanding of the cells that compose the skeletal muscle tissue is limited and molecular definitions of the principal cell types are lacking. Using a combined approach of single-cell RNA-sequencing and mass cytometry, we precisely mapped the different cell types in adult skeletal muscle tissue and highlighted previously overlooked populations. We identified known functional populations, characterized their gene signatures, and determined key markers. Among the ten main cell populations present in skeletal muscle, we found an unexpected complexity in the interstitial compartment and identified two new cell populations. One express the transcription factor Scleraxis and generate tenocyte-like cells. The second express smooth muscle and mesenchymal cell markers (SMMCs). While distinct from MuSCs, SMMCs are endowed with myogenic potential and promote MuSC engraftment following transplantation. Our high-dimensional single-cell atlas uncovers principles of an adult tissue composition and can be exploited to reveal unknown cellular sub-fractions that contribute to tissue regeneration.

## Main Text

Skeletal muscle regeneration is of clinical importance to muscular dystrophies and muscle injuries following trauma. The remarkable regenerative capacity of this tissue arises from a population of resident muscle satellite (stem) cells (*1*) (MuSCs) that express the transcription factor Pax7 and are essential for muscle tissue repair (*2–4*). Other cell types with regenerative potential have also been identified by lineage tracing such as progenitor cells expressing the Twist2 (Tw2) transcription factor (*5*) and pericyte-like cells expressing the Alkaline Phophatase (Alpl) enzyme (*6*). However, these auxiliary muscle progenitor cells appear to depend on the presence of the MuSC population during acute-injury induced muscle regeneration(*2–4*). In most tissues, stem cells interact with their microenvironment and receive signals from neighboring cells to establish and maintain their properties. Likewise, muscle-resident cells such as fibro-adipogenic progenitors (FAPs) (*7,8*), macrophages (*9*) and/or endothelial cells (*10*) are vital components of the niche regulating MuSC function during regeneration. Considering the medical importance of skeletal muscle regeneration and the lack of a molecular understanding of its cellular composition, we set out to achieve a system-wide single-cell profiling of the mononuclear cell population in skeletal muscle to characterize the cellular diversity of this tissue. To do so, we performed single-cell RNA-sequencing (scRNA-seq) (*11*) and single-cell Mass Cytometry (CyTOF) (*12*) using mononuclear cell suspensions of mouse hindlimb muscles (*13*) (Fig. 1A). After normalization and quality control of scRNA-seq profiling, we obtained 6518 cells with an average of 1,158 genes expressed per cell (79,123 mean reads per cell for a total of 18,326 genes detected). We assayed approximately 280,000 cells by CyTOF using a panel of 26 markers that are expressed in various muscle-resident cell types (Supp. Table1). The scRNA-seq data was analysed using the R package Seurat (*14*) with unsupervised graph clustering (*15*) and 10 major cell clusters were identified. We assigned identities based on expression of previously established markers and visualized the data using t-distributed stochastic neighbourhood embedding (*16*) (t-SNE; Fig. 1B, left). We applied standard manual gating to the CyTOF data to identify known muscle-resident cell populations (e.g. FAPs, MuSCs, macrophages and neutrophils – Supp. Fig. 1A). In uninjured muscles, the percentages of known populations relative to the number of mononucleated cells are consistent with findings from previous studies(*17,18*) (Supp. Fig. 1B). To identify distinct cell clusters based on protein expression, we used the viSNE algorithm (*19*) to reduce the multi-dimensional CyTOF dataset to bi-dimensional plots while maintaining the local structure of the data. Each “island” on the graph represents a homogenous population composed by cells sharing similar characteristics (Fig. 1B, right). The plots separated muscle-resident cells into three main groups: blood cells expressing CD45, endothelial cells marked by CD31, and lineage-negative (Lin-) cells containing the highest cellular diversity. The majority of canonical markers were restricted to their specific “island”, confirming the robustness of our approach; only a few markers displayed some promiscuity possibly suggesting a higher degree of similarity between populations (e.g. SCA1 expressed in both FAPs and endothelial cells, CD34 being expressed in MuSCs but also FAPs and endothelial cells) (Fig. 1C and Fig. 1D).

**Fig. 1:**
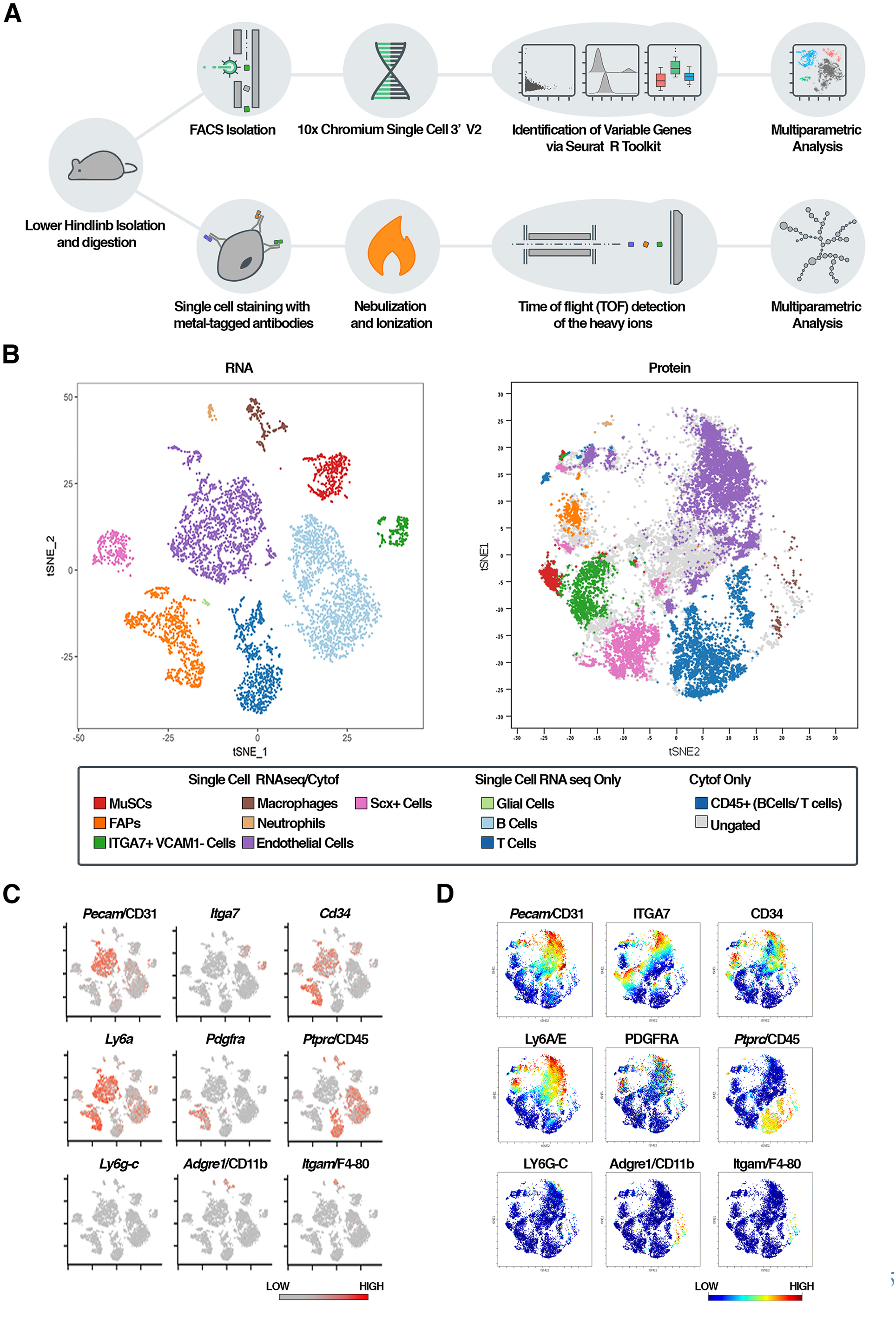
Single cell analysis of skeletal muscle identifies major cell types. **A)** Diagram illustrating the experimental workflow for single cell analyses with 10X Genomics and CyTOF2 platforms. **B)** t-stochastic neighbor embedding (tSNE) plots showing the distribution of main muscle resident populations using both scRNA-seq (left) and CyTOF (right) datasets; each dot represents a single cell. Cells with similar expression profiles were clustered into 10 different groups, as indicated by the different colours. Each cluster was annotated based on the expression of marker genes in the different major cell types. tSNE plots of the scRNA-seq **(C)** and CyTOF **(D)** datasets. Each graph shows the expression pattern of a selected canonical marker used to identify the different populations. Cells are color-coded according to the intensity of the marker shown (representative image of three independent biological replicates). Both names are reported where gene name differs from protein name.

High-dimensional technologies capitalizing on the simultaneous acquisition of multiple parameters allow accurate quantification and identification of the main populations (e.g. MuSCs, FAPs) as well as less abundant cell types (e.g. macrophages, neutrophils) within the tissue. We assigned cluster identity using the scRNA-seq data expression values for specific literature-defined markers (Fig. 2A). Analysis of the differentially expressed genes between the clusters provided molecular definitions for ten main cell populations (Supp. Fig. 1C). Interestingly, while a large number of these markers have already been reported, markers with the highest power of discrimination had been previously overlooked. For example, Asb5 and Cd82 expression levels were well above Vcam1 in MuSCs.

**Fig. 2:**
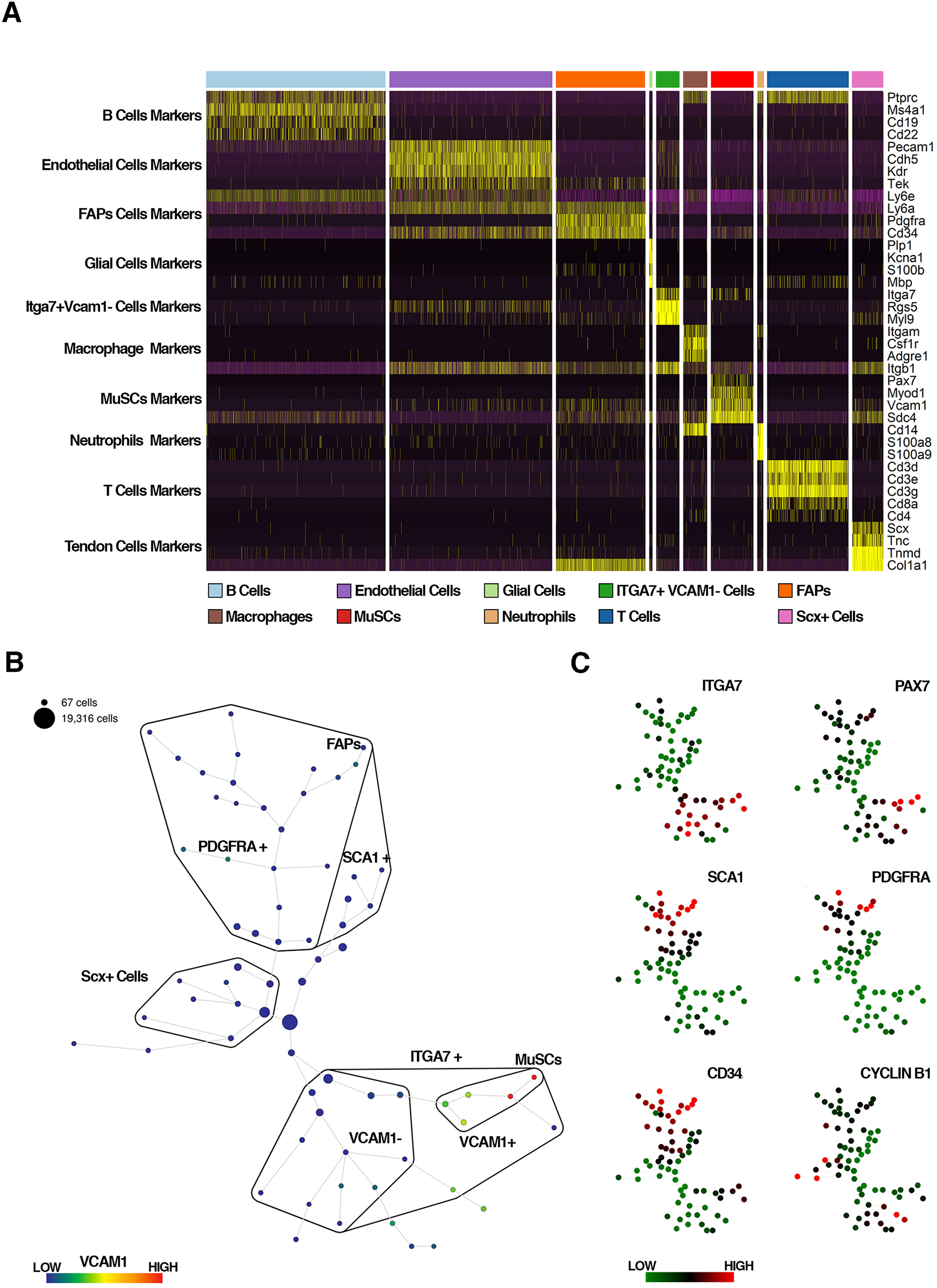
Characterization and hierarchy of muscle cell populations. **A)** RNA Expression heatmap for given cell populations (column) and genes (row), sorted by clusters. Canonical markers used to identify each cluster are plotted (or most variable genes per cluster in case markers were not already present in literature). **B)** SPADE diagram of muscle CD45- CD31- CD11b- (lin-) cells. Nodes of the diagram in the MuSC lineage tree are color-coded according to the relative intensity of expression of VCAM1. Size is proportional to the number of cells assigned to the node. **C)** Selected markers expression levels are overlaid on SPADE diagrams. Each node has the same size and are color-coded according to the intensity of the marker shown. Images are representatives of three independent biological replicates.

In addition, our clustering analysis revealed two novel fractions that have not been described previously. The first cell population expressed Itga7 but not Vcam1, distinguishing them from MuSCs, and the second expressed tenocyte markers and the tendon-specific transcription factor Scleraxis (Scx) (*20*). To understand the relationship of these two cells types with the muscle-resident progenitor cells in the CD45- CD31- (lin-) fraction, we analysed the proteomics dataset with spanning-tree progression analysis of density-normalized events (SPADE) (*27*), which groups similar cells into a defined number of clusters. As expected, the spanning-tree algorithm distinguished different nodes within the VCAM1+/ITGA7+ gate suggesting, as recently reported, a certain degree of cell heterogeneity in MuSCs (*22*). We then observed that in the ITGA7+ fraction, a large number of nodes were negative for most of the MuSC markers (Fig. 2B and Fig. 2C), validating the presence of additional entities. Tendon-like Scx+ cells identified by scRNA-seq, while being in the lin-fraction, were negative for most CYTOF panel antibodies, and located in the continuum between MuSCs and FAPs.

To confirm the existence of muscle-resident cells expressing tendon markers, we analysed cryosections of *Tibialis anterior* (TA) muscles from *Tg:Scx-GFP* reporter mice (*23*). We found that while Scx-GFP was expressed in all developing tendons and ligaments, it also marked interstitial cells outside the myofibres in adult muscle (Fig. 3A). Immunolocalization of the tendon cell markers TNMD (*24*) and COMP (*25*) also identified cells in the muscle interstitium (Fig. 3A). We next isolated muscle-resident tendon-like cells by fluorescence-activated cell sorting (FACS) from *Tg:Scx-GFP* mouse hindlimb muscles. As expected from the CyTOF data, the GFP+ cells were negative for CD31, CD45, SCA1 and VCAM1 expression (Fig. 3B). Sorted cells proliferated *in vitro* and gave rise to flat and elongated cells (Fig. 3C). These cultured cells maintained expression of the *Scx-GFP* transgene, expressed the tendon markers (*26*) TNMD, THBS4, EGR1 and COLA1 and did not express the muscle-specific transcription factor MYOD1 (Fig. 3D). To evaluate if the tendon-like cells could engraft and differentiate *in vivo*, we transplanted 2 × 10^4^ freshly isolated Scx-GFP+ cells into TA muscles of immunodeficient mice (*27*). Three weeks following transplantation, engrafted cells were found in the muscle interstitium, embedded in collagen matrix, suggesting that they may contribute to extra-cellular matrix remodelling during muscle tissue repair (Fig. 3E). Taken together, our experiments demonstrate the existence of a population of interstitial tenocytes in adult skeletal muscle.

**Fig. 3:**
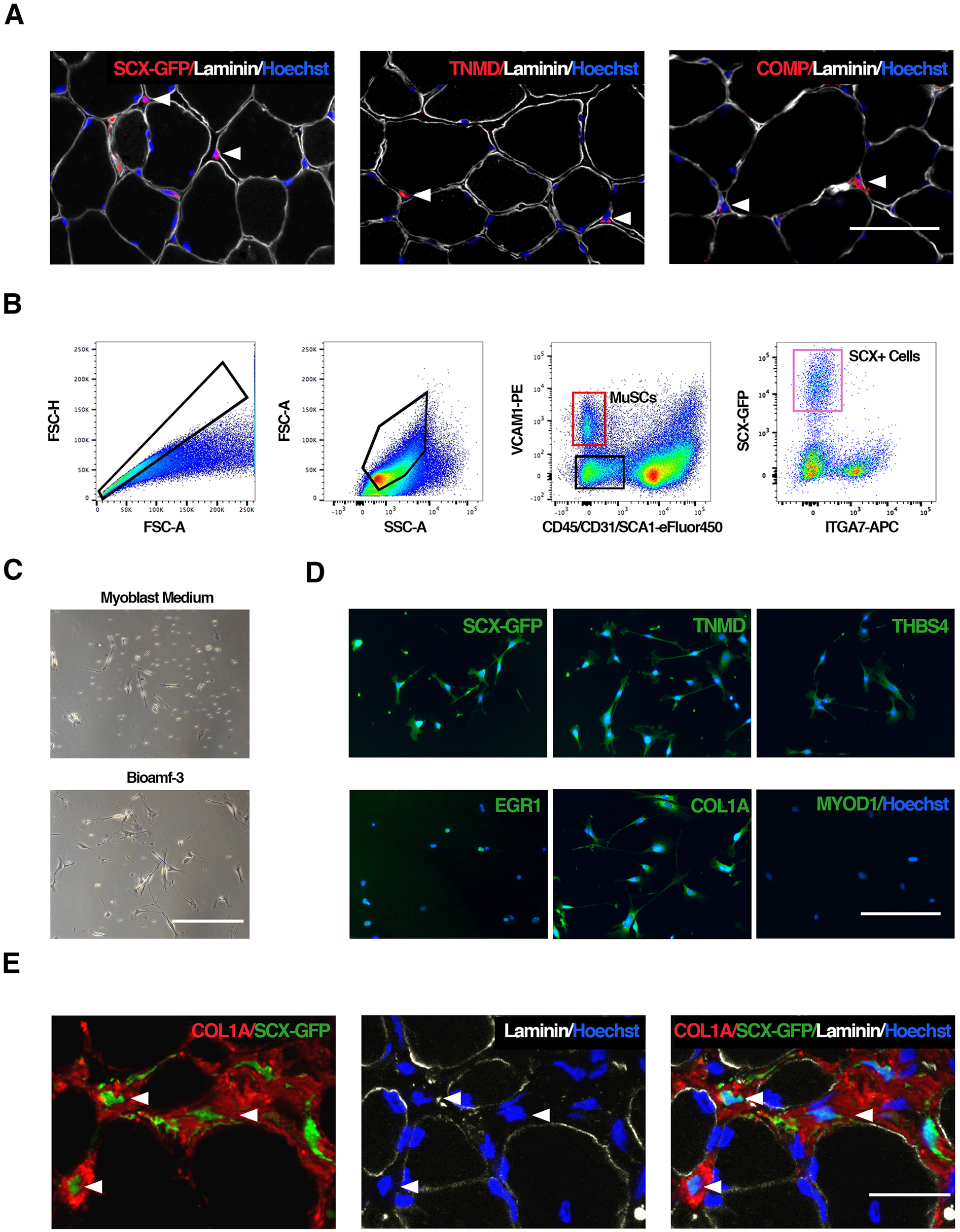
Muscle-resident tenocyte-like cells. **A)** Immunofluorescent stainings of adult (2.5 months old) *Tg:Scx-GFP* mouse Tibialis Anterior (TA) muscle cryosections for indicated markers (scale bar: 100μm). **B)** Flow cytometry analysis of muscle mononuclear cells extracted from *Tg:Scx-GFP* mouse muscles. Representative FACS plots from three different experiments are shown. **C)** Brightfield representative images of sorted Scx-GFP+ cells kept in culture for 6 days (scale bar: 100μm) (N=3). **D)** Immunofluorescent stainings of cultured Scx-GFP+ cells for indicated markers (scale bar: 200μm). **E)** Freshly isolated Scx-GFP+ cells were transplanted into regenerating TA muscles of immunodeficient mice. Immunofluorescent stainings for indicated markers of recipient muscles cryosections, sampled 4 weeks after transplantation (scale bar: 30μm) (N=3). Nuclei are counterstained with Hoechst.

We next prospectively isolated lin-ITGA7+VCAM1-cells by FACS (Fig. 4A) and plated them *in vitro*. When cultured in standard primary myoblast growth medium most of these cells failed to attach and could not proliferate. Out of a panel of different media that were tested, we found that Bioamf3 (Corionic Villli Media), albeit not optimal, enabled the cells to adhere and grow in small clones (Fig. 4B). Once established, the clones maintained a sustained cell growth for several passages. Surprisingly, the *in vitro* descendants of sorted lin-ITGA7+VCAM1-cells activated the expression of the skeletal muscle progenitor marker CXCR4 and of MYOD1 but not PAX7 (Fig. 4C and 4D). Clonogenicity of the sorted cells was found to be 1 in 209 with approximately 30% of the colonies being myogenic (Supp. Fig 2A). When induced to differentiate in low serum conditions, the cells fused together into multinucleated myotubes and expressed Creatine Kinase, Muscle (Ckm) (Fig. 4D) and Myosin Heavy Chains (MyHC) (Fig. 4E). We also tested their ability to differentiate along several cell lineages (adiopogenic, fibrogenic and myogenic) in conditioning media. These lin-ITGA7+VCAM1-cells gave rise to myotubes in each condition tested, substantially failing to display any sign of multipotency (Fig. 4E and Supp. Fig. 2B). Further evaluation of the myogenic differentiation process in lin-ITGA7+VCAM1-descendants showed that the resulting myotubes expressed only MYOSIN-IIX proteins and Myh1 transcript (Fig. 4F), suggesting they may be involved specifically in the maintenance or regeneration of fast glycolytic type IIx muscle fibers *in vivo* (*28*).

**Fig. 4:**
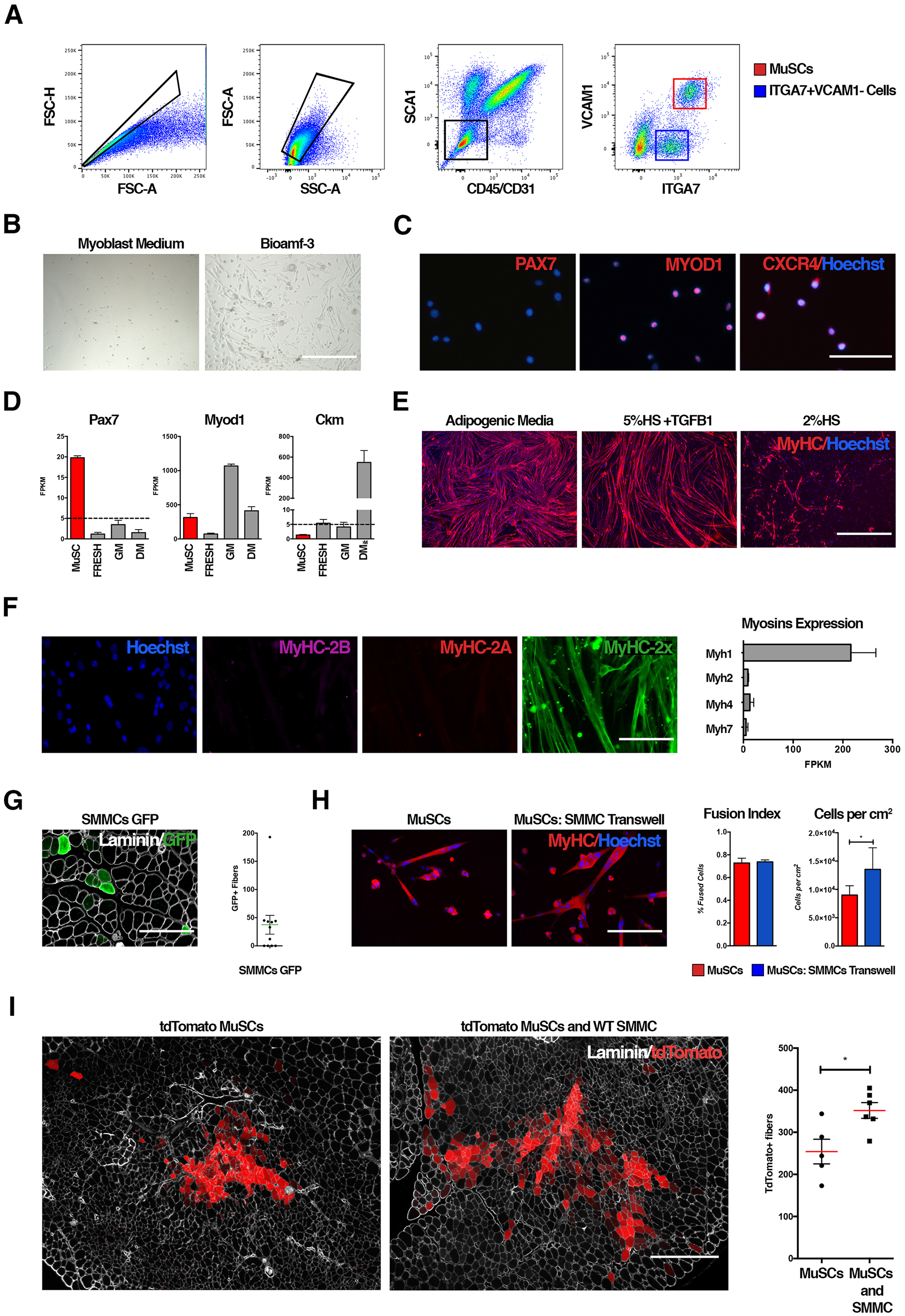
Isolation and characterization of SMMCs. **A)** Flow cytometry analysis of muscle mononuclear cells extracted from adult WT mouse muscles. Representative image from three independent experiments is shown. **B)** Brightfield representative image of lin-ITGA7+VCAM1-cells in culture. Freshly isolated cells were plated in Myoblast Media or Bioamf-3 and kept in culture for 6 days (scale bar: 1000μm) (N=3). **C)** Immunofluorescent stainings of cultured lin-ITGA7+VCAM1-cells for indicated markers (scale bar: 200μm) (N=3). **D)** FPKM (Fragments Per Kilobase of transcript per Million mapped reads) of Pax7, Myod1 and Ckm RNA-seq in quiescent MuSCs and in freshly isolated, growing (Bioamf-3) and differentiated (5%HS) lin-ITGA7+VCAM1-cells. E) Sorted cells differentiation potential (scale bar: 1000μm, N=3). **F)** Immunofluorescent stainings (left) and RNA quantification (right) of different Myosins Heavy Chains (MyHC) isoforms in differentiated cells (left – scale bar: 100μm) (N=3). FPKM of MyHC transcripts in differentiated cells are shown. **G)** Freshly isolated GFP+ SMMCs were transplanted into regenerating TA muscles of immunodeficient mice. Immunofluorescent stainings for GFP (green) and Laminin (white) proteins (left) of recipient muscles cryosections, sampled 3 weeks after transplantation and quantification of GFP+ fibers percentage (right) (N=11) (scale bar: 200μm). Percentage refers to the average of three independent experimental conditions (See also Supp. Table 3). **H)** Transwell co-culture of SMMCs and primary myoblasts derived from MuSCs (N=5). Myoblasts were immunostained for pan-MyHC (red); (scale bar: 200μm). Fusion Index and number of cells per cm^2^ were calculated. (*) *P* < 0.05, Paired two-tailed T-Test (*N* = 5). **I)** Freshly isolated unlabeled SMMCs and tdTomato-labeled MuSCs were co-transplanted into regenerating TA muscles of immunodeficient mice. Immunofluorescent staining for Laminin proteins of recipient muscles cryosections, sampled 4 weeks after transplantation (left). TdTomato-positive fibers are shown in red (scale bar: 400μm). Co-transplantation with SMMCs enhances MuSC engraftment (right). (*) *P* < 0.05, Unpaired two-tailed T-Test with equal variance (MuSC, N=5; MuSC+SMMCs, N=6). Values in bar graph are presented as average ± SEM. Nuclei are counterstained with Hoechst.

To determine if lin-ITGA7+VCAM1-cells are related to MuSCs, we analysed the distribution of CD34 protein and the Pax7-nGFP reporter gene (*29*) in MuSCs and the lin-ITGA7+VCAM1-population by FACS (Supp. Fig 3A – Left). While most MuSCs were positive for both marker genes, lin-ITGA7+VCAM1-cells were negative, indicating that they do not comprise a subpopulation of MuSCs. (Supp. Fig. 3A – Right). These results were validated by quantitative polymerase chain reaction (qPCR) analysis on freshly isolated lin-ITGA7+VCAM1-cells (Supp. Fig. 3B). Of note, the absence of Pdgfra expression by freshly isolated lin-ITGA7+VCAM1-cells further confirmed that they were not FAPs (7,8) nor Tw2+ cells(5). We next labelled MuSCs by inducing Cre mediated recombination of *R26^LSL-tdTomato^* reporter (*30*) in *Pax7^CreERT2^* mice (*3*). We injured hindlimb muscles of tamoxifen-treated mice and analysed this population by FACS after two months when the damaged muscle had completed regeneration (Supp. Fig 4A). While most of MuSCs were tdTomato positive (93.8%), virtually all of the lin-ITGA7+VCAM1-cells (99.1%) were excluded from the tdTomato gate following tissue repair. These observations demonstrate that lin-ITGA7+VCAM1-cells are not activated MuSCs, and that they do not originate from this population in the adult (Supp. Fig. 4B).

Next, we performed bulk RNA-seq on freshly isolated lin-ITGA7+VCAM1-cells and MuSCs and analysed the datasets together with the recently published RNA-seq of Tw2+ cells(*5*) (GEO:GSE84379). Principal component analysis (PCA) of the transcriptional profiles revealed that the three populations were distinct (Supp. Fig. 5A). Gene ontology analysis revealed the most enriched categories in lin-ITGA7+VCAM1-cells, when compared with MuSCs, were mostly related to extracellular space and extracellular matrix, advocating for an interstitial location of these cells within the tissue (Supp. Fig. 5B). Furthermore, the lin-ITGA7+VCAM1-population robustly expressed mesenchymal and smooth muscle cell markers, as well as *Cspg4* (also known as NG2, expressed by interstitial and perivascular cells) (*37*), but the complete absence of *Alpl* expression distinguished them from pericytes (*6*) (Supp. Fig 5C). Therefore, we designated the lin-ITGA7+VCAM1-cells as Smooth Muscle/Mesenchymal Cells (SMMCs).

We next sought to determine whether SMMCs display myogenic potential *in vivo* by isolating 2 × 10^4^ of these cells by FACS from mice ubiquitously expressing GFP and immediately injecting them into pre-injured TA muscles of immunodeficient mice. Three weeks following injection transplanted SMMCs contributed to the formation of GFP+ myofibers (Fig. 4G). However, the number of myofibers and the overall extent of the contribution to the muscle were relatively low compared to MuSC transplantations reported previously (32–34). Co-culture of SMMCs and MuSCs demonstrated that descendants of both cell types generated chimeric myotubes, suggesting that the two populations could interact during muscle regeneration (Supp. Fig. 6). We then used a cell culture insert system to evaluate the effect of secreted signals from SMMCs on MuSCs in differentiation conditions. Compared with MuSCs plated alone, sharing medium with SMMCs promoted myoblast survival while not impeding muscle cell fusion *in vitro* (Fig. 4H). To investigate the possible functional interaction between MuSCs and SMMCs *in vivo*, we performed co-transplantation experiments. Permanently labelled MuSCs from *Pax7^CreERT2^;R26^L.SL-tdTomato^* mice were transplanted alone or in combination with unlabelled SMMCs into pre-injured TA muscles of immunodeficient mice. Quantification of the number of tdTomato+ myofibers in transplanted muscles 4 weeks after injection revealed that cotransplantation consistently give rise to a higher number of myofibers (Fig. 4I), suggesting that SMMCs synergize with MuSCs *in vivo* and play a role as helper cells supporting MuSCs during muscle regeneration.

In summary, using a novel combined single-cell proteomics and transcriptomics approach we have determined the cellular composition and the molecular signature of each cell type in adult mouse skeletal muscle. The blueprint presented here yields crucial insights into muscle-resident cell type identities and will serve as resource for the field. To prove the strength of this approach, we selected two new previously unidentified populations from our screening and characterized them both *in vivo* and *in vitro*. This analysis led to the identification of a population of muscle resident tenocytes-like cells as well as a myogenic subset of SMMCs that did not exhibit multipotency. SMMCs can fuse with MuSCs and promote muscle regeneration through paracrine mechanisms, but appear to be distinct from other non-MuSCs muscle progenitors(*5*),(*6*). Our data clearly demonstrates that this strategy can be successfully applied to deconstruct and identify diverse cell types in other tissues and organs both in health and disease.

The accompanying datasets the will also serve as reference for future single-cell profiling in the context of skeletal muscle aging or in disease such as muscular dystrophies. Single-cell resolution mapping in a pathological background will likely lead to the identification of novel biomarkers and facilitate the identification of dysfunctional cellular subpopulations, their relative proportions and molecular signatures, which lay the foundations for the development of new therapeutic strategies for neuromuscular diseases.

## Acknowledgments

We thank the CyPS Facility for technical support. We thank C. Combadière for providing ubiquitous GFP mice and G. Comai for providing Scx-GFP mice. We thank G. Butler-Browne, D. Sassoon, R. Mounier and C. Trollet for commenting the draft manuscript.

## Author contributions

L.G., F.L.G and T.H.C conceived and designed the study. L.G. performed most of the experiments with the assistance of M.M.S. at the late stage of the project. F.L.G performed cell culture experiments. G.H. performed the single-cell RNA-seq experiments. R.W. performed the data analysis of the RNA-seq experiments. H.S. and S.T. provided mouse models. E.N. performed transplantation experiments. J.L. performed bulk RNA-seq. L.G. and F.L.G. wrote the manuscript with substantial input from S.T. and T.H.C. All authors reviewed the manuscript.

**Supplementary Materials**

Materials and Methods

Fig. S1: CyTOF gating strategy and top 20 discriminating genes.

Fig. S2: Limiting Dilution assay and myogenic frequency for SMMCs.

Fig. S4: No Pax7-TdTomato+ progeny in SMMC gate.

Fig. S5: RNA-seq analysis of Tw2+ cells, MuSCs and SMMCs.

Fig. S6: Co-culture experiment of SMMCs and MuSCs *in vitro*.

Sup. Table 1: Complete list of metal conjugated antibodies used in CyTOF experiments.

Sup. Table 2. Frequency of SCX+ and SMMCs

Sup. Table 3: Number of GFP+ fibers in each transplant condition tested for SMMCs.

Sup. Table 4: Lists of all transcripts used to assign cell population identity, primers, antibodies and publicly available datasets used in the paper.

References (35–44)

## Materials and Methods

### Animals

Animals were handled according to European Community guidelines. Experimental animal protocols were performed in accordance with the guidelines of the French Veterinary Department and approved by the Sorbonne Université Ethical Committee for Animal Experimentation. In this study 10-weeks-old WT C57/BL6 mice were used. *Tg:Pax7-nGFP, Pax7^CreERT2^;R26^LSL-tdTomato^* and *Tg:Scx-GFP* mice were crossed and maintained on a F1 C57/BL6:DBA2 background and genotyped by PCR. *Rag2^-/-^;Il2rb^-/-^* mice and *Rag2^-/-^;Il2rb^-/-^*;*Mdx*^DMD-/-^ were provided by Dr. Vincent Mouly (Myology Institute – Paris). *Tg:UBI-GFP* and *Tg:H2B-eGFP* mice were kindly provided as gifts from Dr. Christophe Combadière (Centre d’Immunologie et des Maladies Infectieuses – Paris) and Dr. Bruno Cadot (Myology Institute – Paris) respectively. Cardiotoxin injections and Tamoxifen treatments were performed as previously described (35).

### Single Cell Preparation

Single cell suspension from hindlimb muscles was prepared by mechanical and enzymatic dissociation as previously described with minor modification. Briefly, hindlimb muscles were dissected and minced into a fragmented muscle suspension. The muscle suspension was firstly digested with Collagenase II (1000U/ml) in Ham’s F10 containing 10% horse serum for 90 min, followed by further digestion with Collagenase II (1000U/ml) and Dispase (11U/ml) for 30 min. The digested suspension was then triturated and washed to yield a mononuclear cell suspension. To improve dissociation cells were passed 10 times in a 20-gauge needle syringe and then filtered with a 35 μm cell strainer.

### Single-cell RNA sequencing and analysis

Single cell suspension was then subjected to BD Influx cell sorter to isolate single cells with debris and doublets excluded. Cells were pelleted and washed with 1X PBS with 0.04% BSA twice so as to remove ambient RNA as well as minimize cell aggregation. Cells were filtered using a 70μm cell strainer to remove any remaining cell debris and large clumps. Finally, cell concentration was determined using hemocytometer and adjusted to obtain the target concentration for the following 10x Chromium system chip loading. Cells were then loaded into the 10x Chromium system and went through the Single Cell 3’ Reagent Kits v2 as per the manufacturer’s protocol. Following library preparation and quantitation, libraries were sequenced on Illumina NextSeq 500 platform using High Output Kit v2 (300 cycles) with customized sequencing run parameters (Read 1 – 30 cycles, i7 index – 8 cycles, Read 2 – 151 cycles). Raw sequencing data was processed by Cell Ranger 2.1.0 (10X Genomics) to generate a gene-cell expression matrix. Further data analysis was carried out in R version 3.4.3 using Seurat version 2.2. “Cells” that fit any of the following criteria were filtered out: <350 or >9000 UMIs, <350 or >4000 expressed genes, or >20% UMIs mapped to mitochondria. Finally, we obtained 6,518 cells that passed quality control with an average of 1158 gene expressed per cell (79123 mean reads per cell for a total of 18326 genes detected). The top 2000 variably expressed genes (with highest dispersion across single cells) were subjected to PCA for dimensionality reduction. The top 20 significant PCs that explained more variability by chance were selected for downstream graph-based clustering and t-SNE visualization. Further, we manually assigned cell population identity based on cell type specific markers and identified 10 different cell types (see Supplementary Table 4 for the list of markers used). After clustering and cell population identification, the most highly differentially expressed genes or cluster putative markers were identified by likelihood-ratio test, as implemented in Seurat.

### Bulk RNA-seq

The mouse genome and its annotations were obtained from UCSC (mm 10) via Illumina’s iGenomes web site. For the study were used three SMMCs biological conditions (freshly isolated, proliferating, and differentiating; N=3 each) and freshly isolated MuSC (QSC) (N=2). Additionally Twist2 datasets from Liu et al. ^9^ (Twist-2 positive and Twist-2 48h samples) were downloaded from GEO website; see supplementary Table 4 for a complete list of the dataset used for comparison. All experiments were performed on a computing cluster running CentOS version 6.6. All softwares were executed within a Conda 4.3.21 environment. Datasets were processed according to most of the steps outlined in Pertea et al. (36). Whenever possible, the gene annotation file was also used. First, the genome index for mm10 was created with Hisat2 (version 2.1.0) using splice-sites and exon information from the gene annotation file according to [6]. Alignment to the genome was performed using Hisat2 (37). A sorted BAM file was created using SAMtools (version 1.3.1) (38). Then, StringTie (version 1.3.3) (39) was used with the -e option to quantify the expression level of genes and transcripts. The -e option limits the processing of read alignments to only those that match the reference transcripts [https://ccb.jhu.edu/software/stringtie/index.shtml?t=manual]. Transcripts were merged across all samples using the –merge option to Stringtie. Finally, the expression level of transcripts was used to create table counts for Ballgown using the -B option to Stringtie. Ballgown version 2.2.0 was used with R version 3.3.1 (40). After extracting the genes from the Ballgown object, genes whose mean expression level (in FPKM) across all samples was less than 1 were removed. Then, only genes which have an FPKM of at least 5 in at least one sample were retained. These sets of genes were used to create the Principle Component Analysis (PCA) plots with the prcomp () function in R with both centering and scaling set to TRUE.

### Mass Cytometry (CYTOF)

After filtration, single cell preparation was resuspended in Ham’s F10 at the concentration of ~ 3 × 10^6^ cells/ml and treated with IdU at a concentration of 50μm for 25 min at 37°C 5%CO_2_. Then cells were washed with F10 10%HS and resuspended in Staining Buffer (Fludigm). From this step on all washes unless otherwise stated were done in Staining Buffer (SB) Surface antibodies cocktail was added to the cells to reach a final cell concentration of ~ 3 × 10^7^ per ml. Surface staining was performed at 4°C for 30 min. Cells were then washed twice and fixed with 2%PFA for 15 min. After PFA fixation, cells were washed again and left to pre-chill on Ice for 10 min then fixed with cold methanol for 15 min on ice. Cells were washed twice and then stained with intracellular antibodies cocktail for 45 minutes at room temperature at a final cell concentration of ~ 3 × 10^7^ per ml. Cells were then washed twice and incubated overnight with the intercalator buffer (Intercalator 0,1% in Maxpar Fix and Perm – Fluidigm). Cells were then washed twice in SB and once in ddH_2_O water. Cells were diluted in (dd)H_2_O containing beads standards to allow for sample normalization and then run on a CyTOF2 or an Helios Mass cytometer (Fluidigm). All mass cytometry files per experimental condition were normalized together. After removal of cell debris data analysis has been performed on Cytobank (41) platform.

### FACS

After filtration, single cell preparation was resuspended in Ham’s F10 containing 10% horse serum and stained with antibodies for 30 min at 4 °C in the dark (See Supplementary Table 4 for a list of antibodies used). Cells were either analyzed by LSRFortessa or sorted on a BD FACSAria II. Debris and dead cells were excluded by forward scatter and side scatter. Scx-GFP+ cells were isolated from *Tg:Scx-GFP* mice as CD45^neg^ CD31^neg^ SCA1^neg^ VCAM1^neg^ ITGA7^neg^ GFP^pos^. SMMC cells were isolated from C57/BL6 mice as CD45^neg^ CD31^neg^ SCA1^neg^ VCAM1^neg^ ITGA7^pos^.

### Immunostaining

Cryosections were fixed with 4% PFA in PBS (Electron Microscopy Sciences) then incubated with blocking solution (BSA 2.5%, 2% goat serum, 0.1% Triton-100X) for 1h. After the blocking step, cryosections were incubated overnight with primary antibodies at 4°C. For the detection of TdTomato or GFP proteins whole TA muscles were pre-fixed prior to freezing in 2% PFA 0.2% Triton-100X for 2h and incubated in 15% Sucrose in PBS overnight at 4°C. Cells were fixed with 4% PFA, permeabilized with 0.25% Triton X-100, blocked with 4% BSA, and incubated with primary antibodies overnight at 4°C. Detection of primary antibodies was achieved using Alexa Fluor secondary antibody (Invitrogen) at a 1:1000 dilution in PBS. Nuclei were counterstained with Hoechst at 0.5 μg/ml. See Supplementary Table 4 for a list of antibodies used. Oil Red O (Sigma-Aldrich) staining was performed according to manufacturer’s instructions.

### Transplantation

For Scx-GFP+ cells transplantation experiment TA muscles of *Rag2^-/-^; Il2rb^-/-^; Mdx^DMD-/-^* were transplanted with ~ 2 × 10^4^ cells, freshly isolated from *Tg:Scx-GFP* mouse. Recipients TA were injured with CTX 24h before transplantation. Animals were sacrificed 4 weeks after transplantation (N=3). For SMMCs transplant were performed on *Rag2^-/-^;Il2rb^-/-^ and Rag2^-/-^; Il2rb^-/-^; Mdx^DMD-/-^* mice using ~ 2 × 10^4^, freshly isolated from *Tg:UBC-GFP* mice. TA muscles of *Rag2^-/-^;Il2rb^-/-^* mice were injected with SMMCs 72h post CTX injury and animals sacrificed after 3 weeks (N=4). For TAs of *Rag2^-/-^; Il2rb^-/-^; Mdx^DMD-/-^* injection was performed either 48h post irradiation (N=4) or 24h post CTX injury (N=3). In both conditions animals were sacrificed after 4 weeks. For co-transplantiation experiments MuSCs and SMMCs were freshly isolated from *Pax7^CreERT2^;R26^LSL-tdTomato^* mice (previously treated with Tamoxifen to induce CRE mediated recombination). *Rag2^-/-^; Il2rb^-/-^; Mdx^DMD-/-^* mice were then injected with MuSCs (~ 2 × 10^4^ cells) or with MuSC in combination with SMMCs (~ 2 × 10^4^ cells each). Recipients TA were injured with CTX 24h before transplantation. Animals were sacrificed 4 weeks after transplantation (N=5 MuSCs; N=6 MuSC+SMMCs).

### RT-qPCR

RNA was extracted using Direct-zol Miniprep Plus (Zymogen) and reverse transcriptase step was performed using QuantiTect Reverse Transcription Kit (Qiagen). Quantitative PCR was carried out in a LightCycler 96 (Roche) using FastStart Universal SYBR Green Master (Rox) (Roche). Each step has been performed according to manufacturer’s instructions. See Supplementary Table 4 for the list of primers used.

### Cell Culture

MuSCs and primary myoblasts were cultured on collagen-coated dishes. Myoblast Medium consisted of Ham’s F10 medium (Life Technologies) supplemented with 20% FBS (Eurobio), 2ng/ml basic fibroblast growth factor (FGF) (R&D Systems) and 1% penicillin/streptomycin (Pen/Strep) (Life Technologies). SMMCs and SCX-GFP+ cells were cultured on 1 mg/ml Matrigel matrix (Corning Life Sciences) either in Myoblast Medium or in Bioamf-3 complete medium (Biological Industries). To test differentiation potential, freshly sorted SMMCs were allowed to adhere and grow on matrigel-coated plates for 6 days in Bioamf-3. Cells were then tripsinized and plated at a density of ~ 1 × 10^4^ cells per cm^2^. After 24h cells were switched to differentiation conditions. To promote myogenic differentiation cells were exposed to DMEM supplemented with 2% horse serum (Sigma) and 1% Pen/Strep (Life Technologies) for at least 48h. To induce adipogenic differentiation cells were cultured for 3 days in Adipogenic Induction medium followed by Adipogenic Maintenance media, as described previously^11^. For fibrogenic differentiation cells were kept for 6 days in DMEM 5% horse serum (Sigma) supplemented with 5ng/ml TGF-β1 (R&D Systems) and 1% Pen/Strep (Life Technologies). To avoid myotubes detachment matrigel was layered on the cells 48hrs after induction of differentiation (1:2 in DMEM to a final concentration of 0.5mg/ml) (42). Transwell experiments were performed using ~ 2 × 10^4^ freshly isolated SMMCs and primary myoblasts. SMMCs were plated into transwells and left to proliferate in Bioamf-3 for 5 days. Myoblasts were plated at a density of ~ 2 × 10^4^ cells per cm^2^ in GM. After 48h, myoblasts where shifted in 5%HS and left to differentiate in presence or absence of the SMMCs-containing transwell. Myoblasts were fixed after 5 days.

### Limiting Dilution Assay and Myogenic Frequency

To assess clonogenicity, 1 up to 250 cells were sorted into individual 96 multiwell matrigel-coated wells. After three weeks in Bioamf-3 wells were scored for the presence of colonies. A minimum of 30 replicates for at least 3 biological replicates has been tested for each dose (respectively 1, 25, 50, 100, 250 cells per well). Data were analyzed as previously described^12^ using a web application made available by the Walter and Eliza Hall Institute of Medical Research, Melbourne, Australia (http://bioinf.wehi.edu.au/software/elda/index.html) (43). To assess frequency of Myogenic clones 100 cells were plated in replicate wells of a 96 multiwell directly from FACS. The number of cells was selected to obtain a ratio of 0.5 clone forming cells per well to ensure the monoclonality of the derived population. After 10 days, cells where shifted in 5% HS for five days to promote myogenic differentiation. Wells were scored for the presence of colonies and the fraction of myogenic versus non-myogenic was calculated. Only colonies larger than 100 cells were considered to ensure that only those wells that contained a cell able to undergo proliferation would be taken in account.

**Fig. S1:**
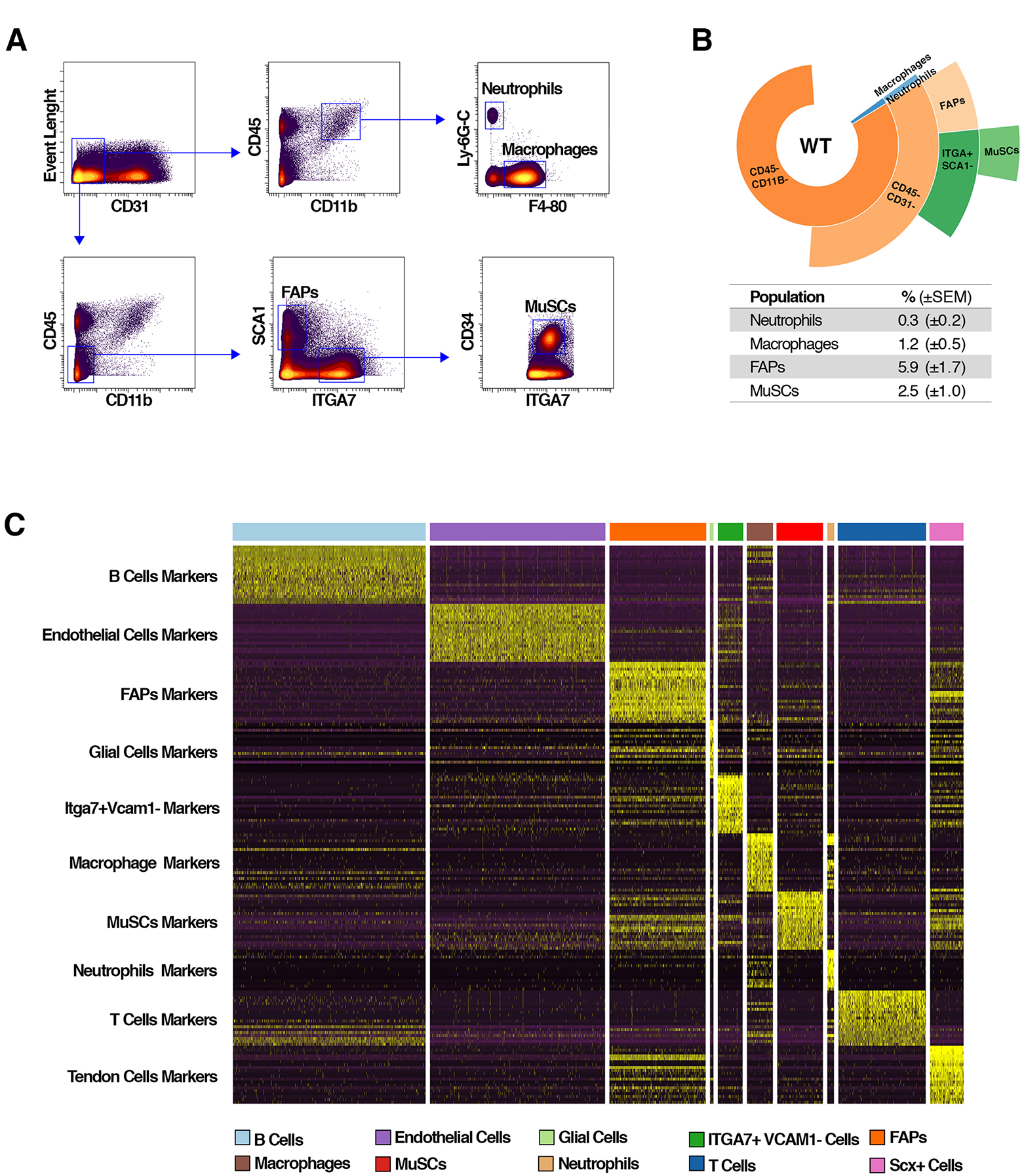
CyTOF gating strategy and top 20 discriminating genes. **A)** CyTOF dot-plot illustrating the gating strategy applied to identify different resident cell populations in the skeletal muscle. (N = 6) **B)** Sunburst plot with the relative percentage of the population identified (N = 3; representative image). **C)** RNA Expression heatmap for given cells (column) and genes (row), sorted by clusters. Top 20 most discriminating markers for each cluster are plotted.

**Fig. S2:**
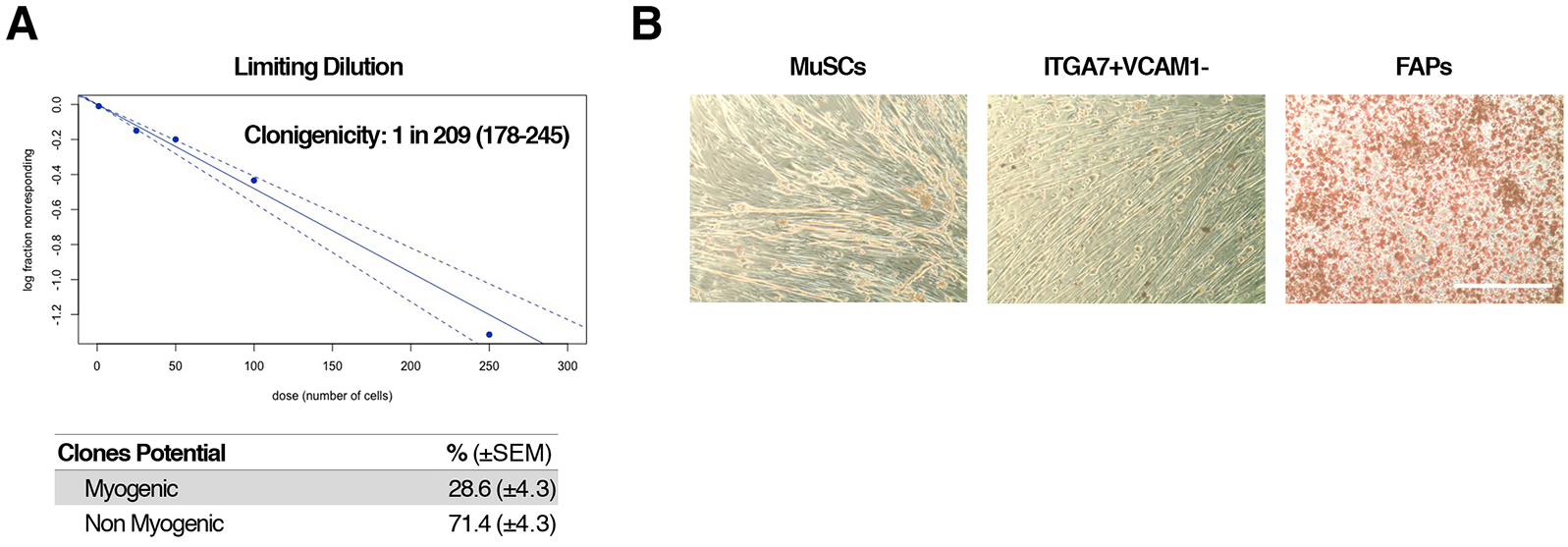
Limiting Dilution assay and myogenic frequency for SMMCs. **A)** Limiting dilution assay (top) and frequency of myogenic clones in SMMCs (bottom) (mean ± SEM.; N=3). **B)** Oil Red O staining of SMMCs, MuSC and FAPs in adipogenic media (scale bar: 200μm).

**Fig. S3:**
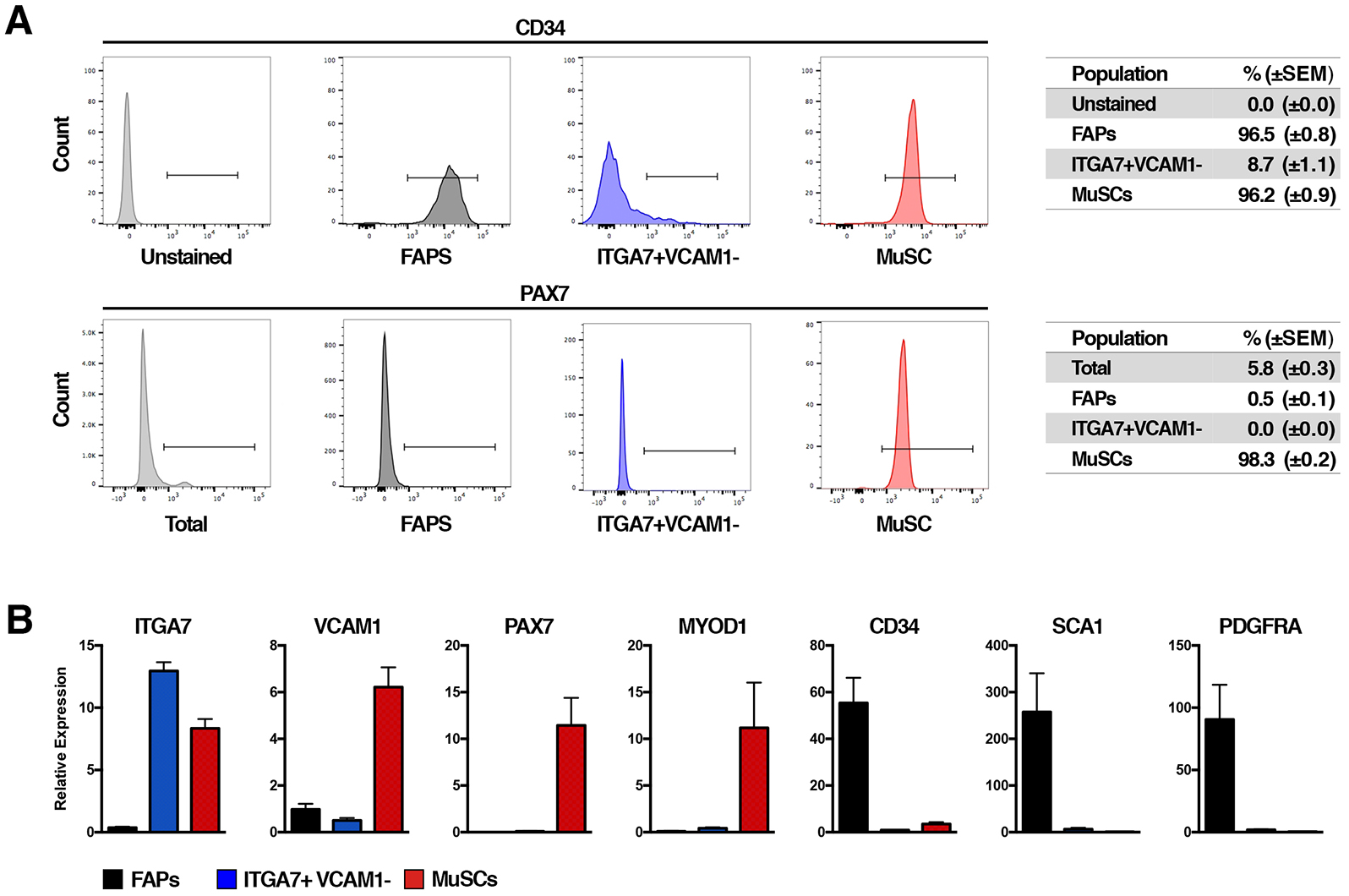
CD34 and PAX7 expression in MuSCs FAPS and SMMCs. **A)** FACS profiles of FAPs, SMMCs and MuSCs. Cells were analyzed for CD34 and Pax7-GFP expression. Table on the right indicates the relative percentage of the population that falls within the positive gate (mean ± SEM; N= 3). **B)** Relative gene expression of selected genes in freshly isolated FAPs, MuSCs and SMMCs (mean ± SEM.; N=3).

**Fig. S4:**
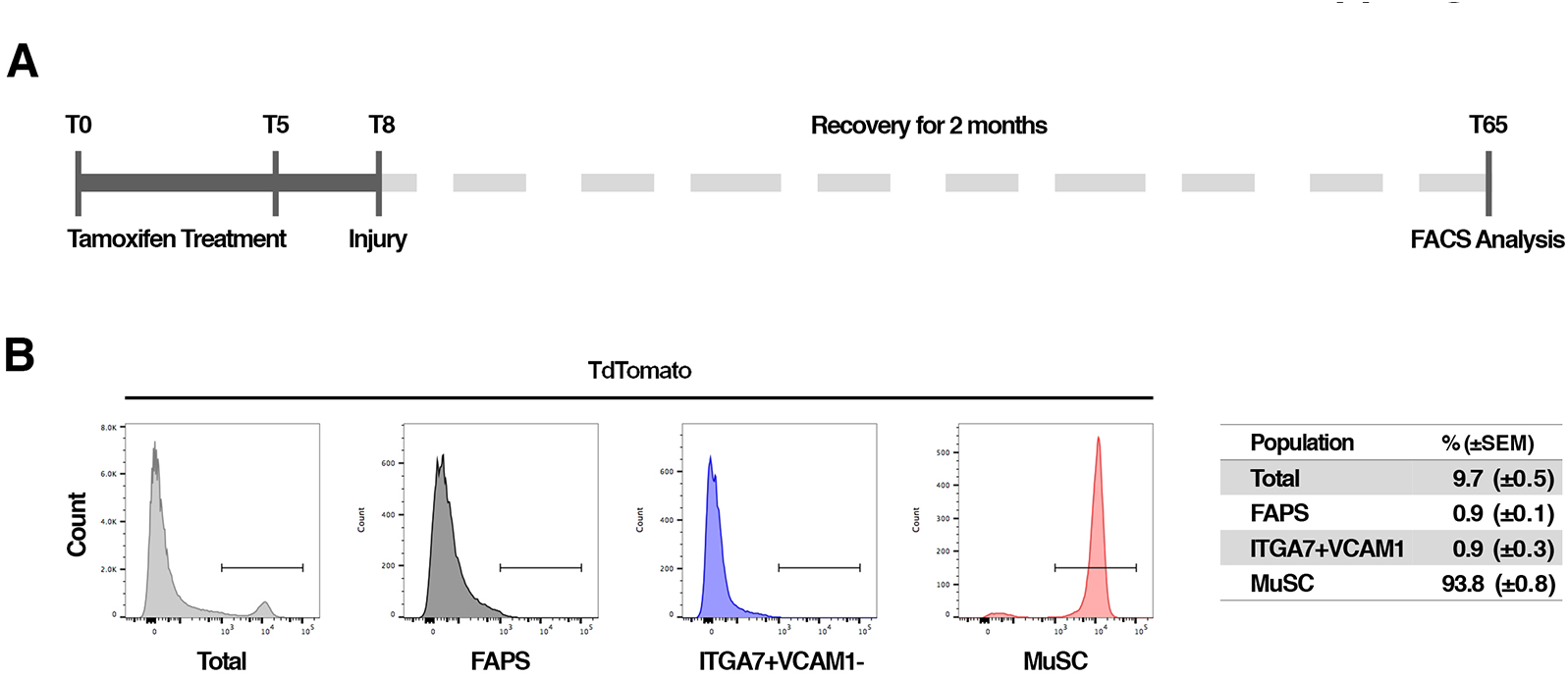
No Pax7-TdTomato+ progeny in SMMC gate. **A)** Experimental outline of *Pax7^CreERT2^;R26^LSL-tdTomato^* injury experiment. Mice were treated with tamoxifen to trigger CRE mediated recombination and permanent labeling of Pax7+ cells. 7 days post tamoxifen treatment, TA muscles were injected with CTX to induce muscle injury. After 2 months animals were sacrificed and scored for the presence of tdTomato-positive cells in the SMMCs gating. **B)** Representative FACS profile of FAPS, SMMCs and MuSCs from experiment described in A. Cells were analyzed for tdTomato expression. Table on the right indicate the relative percentage of the population that falls within the positive gate (mean ± SEM; N=3).

**Fig. S5:**
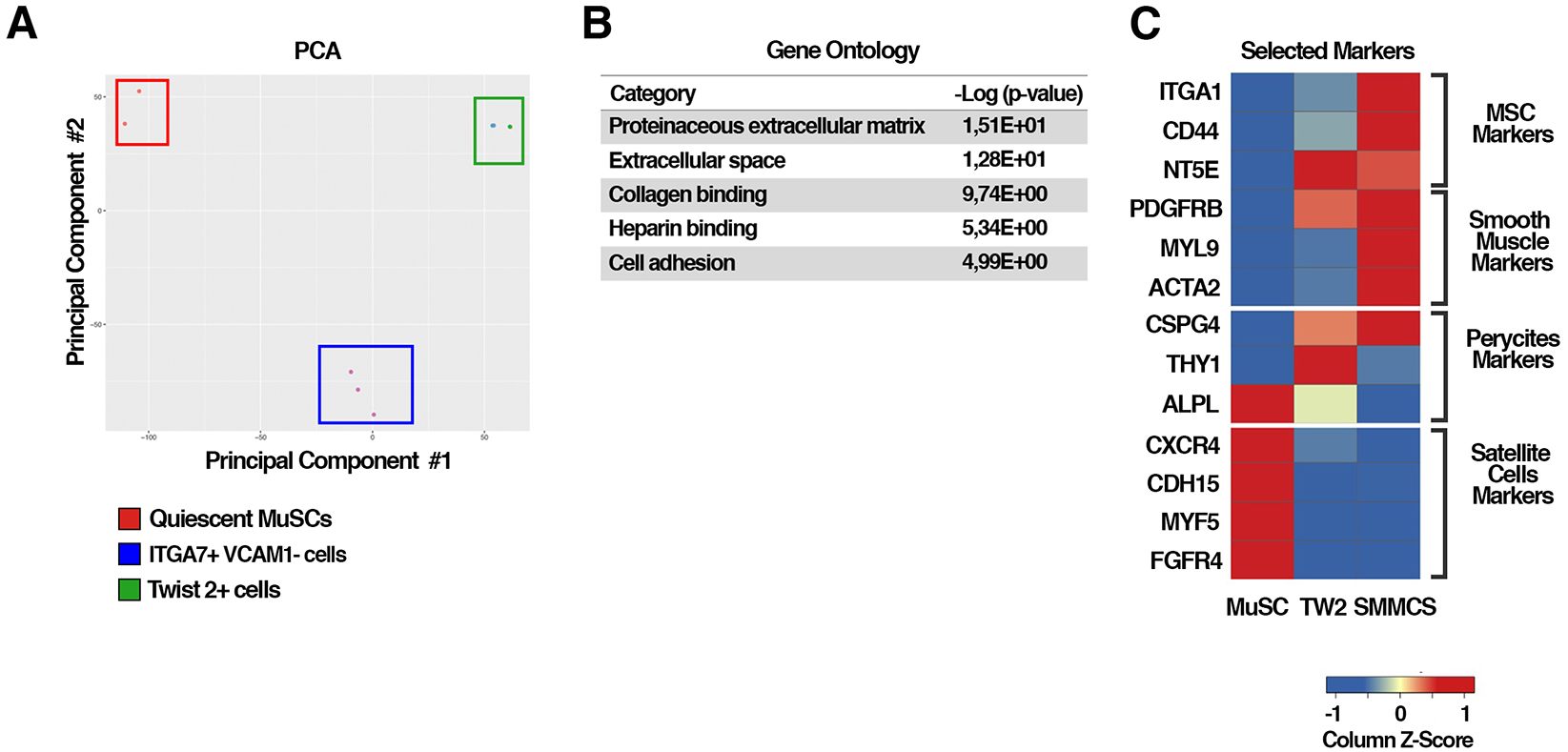
RNA-seq analysis of Tw2+ cells, MuSCs and SMMCs. **A)** Principal component analysis of MuSCs (red box), SMMCs (blue box) and Tw2+ cells (green box). **B)** Gene ontology categories enriched in freshly sorted SMMCs cells identified by DAVID. **C)** Heatmap of MSCs, Smooth Muscle Cells, Pericyte and MuSCs marker genes identified by RNA-seq expressed in freshly isolated quiescent MuSCs, TW2 cells and SMMCs.

**Fig. S6:**
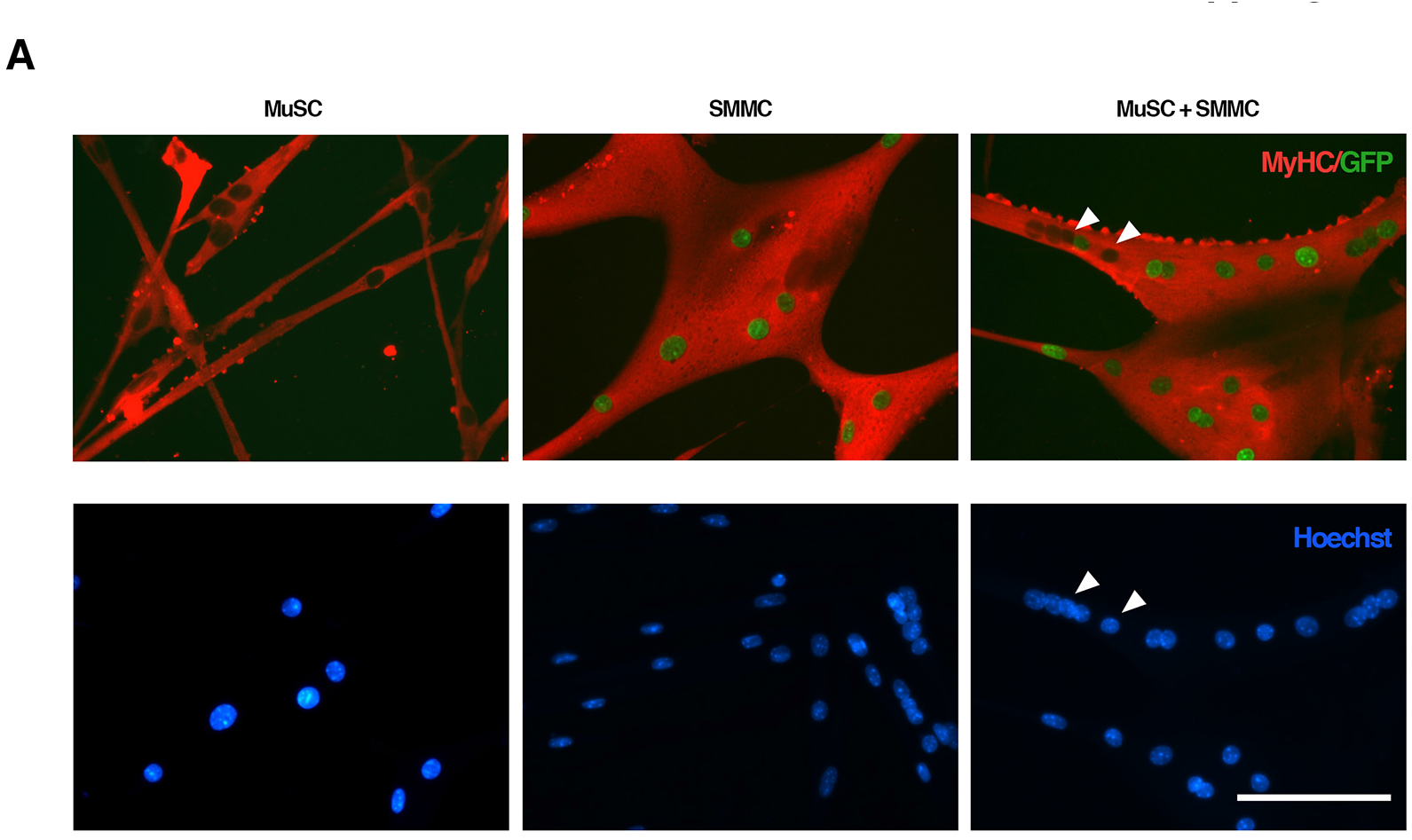
Co-culture experiment of SMMCs and MuSCs *in vitro*. A) SMMCs isolated from GFP-H2B mouse, WT MuSCs or an equal ratio of the two populations were plated at a density of ~ 1 × 10^4^ cells per cm^2^ in Bioamf-3. After 24h, cells were shifted in 5%HS for 3 days and then immunostained for GFP (green) and Embryonic Myosin Heavy chain (red) proteins. Nuclei were counterstained with Hoechst (scale bar: 100μm).

**Table S1.** Complete list of metal conjugated antibodies used in CyTOF experiments.

| Exp1 Fig1 |  |  | Exp2 Fig2 |  |  |
| --- | --- | --- | --- | --- | --- |
| Marker | Metal | Producer | Marker | Metal | Producer |
| Ly6C | Dy162 | Fluidigm | Ly6C | Dy162 | Fluidigm |
| B-Catenin | Sm147 | Fluidigm | B-Catenin | Sm147 | Fluidigm |
| CD64 | Eu151 | Fluidigm | CD64 | Eu151 | Fluidigm |
| CyclinB1 | Eu153 | Fluidigm | CyclinB1 | Eu153 | Fluidigm |
| Tgfb | Sm154 | AbLab | Tgfb | Sm154 | AbLab |
| ITGA7 | Gd160 | AbLab | ITGA7 | Gd160 | AbLab |
| IFNg | Ho165 | AbLab | Active Bcat | Ho165 | Fluidigm |
| CD31 | Er168 | AbLab | CD31 | Er168 | AbLab |
| CD34 | Er170 | Fluidigm | CD34 | Er170 | Fluidigm |
| CD106 | Yb174 | AbLab | CD106 | Yb174 | AbLab |
| pH3 | Lu175 | Fluidigm | pH3 | Lu175 | Fluidigm |
| Ly6G-C | Pr141 | Fluidigm | Ly6G-C | Pr141 | Fluidigm |
| CD45 | Nd145 | AbLab | CD45 | Nd145 | AbLab |
| CD11b | Nd148 | Fluidigm | CD11b | Nd148 | Fluidigm |
| F4-80 | Tb159 | Fluidigm | F4-80 | Tb159 | Fluidigm |
| IL-6 | Er167 | Fluidigm | GFP | Er167 | AbLab |
| Ly6A-E | Tm169 | Fluidigm | Ly6A-E | Tm169 | Fluidigm |
| CD86 | Yb172 | Fluidigm | CD86 | Yb172 | Fluidigm |
| pAkt | Sm152 | Fluidigm | pAkt | Sm152 | Fluidigm |
| TNFa | Nd143 | AbLab | TNFa | Nd143 | AbLab |
| pRb | Nd150 | Fluidigm | pRb | Nd150 | Fluidigm |
| p-p38 | Gd156 | Fluidigm | p-p38 | Gd156 | Fluidigm |
| pStat3 | Gd158 | Fluidigm | pStat3 | Gd158 | Fluidigm |
| PDGFRa | Dy164 | AbLab | PDGFRa | Dy164 | AbLab |
| IL-4 | Er166 | Fluidigm | Idu | l127 | Fluidigm |
| Idu | l127 | Fluidigm | IL-10 | 176Lu | AbLab |
| N=3 |  |  | CD206 | 144Nd | AbLab |
|  |  |  | IFNg | 173Yb | AbLab |
|  |  |  | IL-6 | 139La | AbLab |
|  |  |  | N=3 |  |  |

**Table S2.** Frequency of SCX+ and SMMCs.

| Strain | Mouse Number | % of SCX+ Cells |
| --- | --- | --- |
| Scx-GFP | 1 | 1,5 |
|  | 2 | 1,3 |
|  | 3 | 1,4 |
|  | 4 | 1,7 |

|  | Average | SEM |
| --- | --- | --- |
| SCX+ | 1,48 | 0,09 |

| Strain | Mouse Number | % of SMMCs |
| --- | --- | --- |
| C57BL6 | 1 | 3,9 |
|  | 2 | 3 |
|  | 3 | 3,5 |
|  | 4 | 4,3 |
|  | 5 | 5 |
|  | 6 | 3,7 |

|  | Average | SEM |
| --- | --- | --- |
| SMMSC | 3,90 | 0,28 |

**Table S3.** Number of GFP+ fibers in each transplant condition tested for SMMCs.

| Donor | Host |  |  |
| --- | --- | --- | --- |
| Ub-eGFP | Rag2 | Rag2/DMD |  |
|  | Day3 PI | Irradiated | Day1 PI |
| Mouse 1 | 42 | 43 | 0 |
| Mouse 2 | 33 | 44 | 193 |
| Mouse 3 | 12 | 0 | 44 |
| Mouse 4 | 0 | 0 | \ |
| **Average** | **21,75** | **21,75** | **59,25** |

| Average GFP+ fibers | SEM |
| --- | --- |
| 34,25 | 16,23 |

**Table S4.**
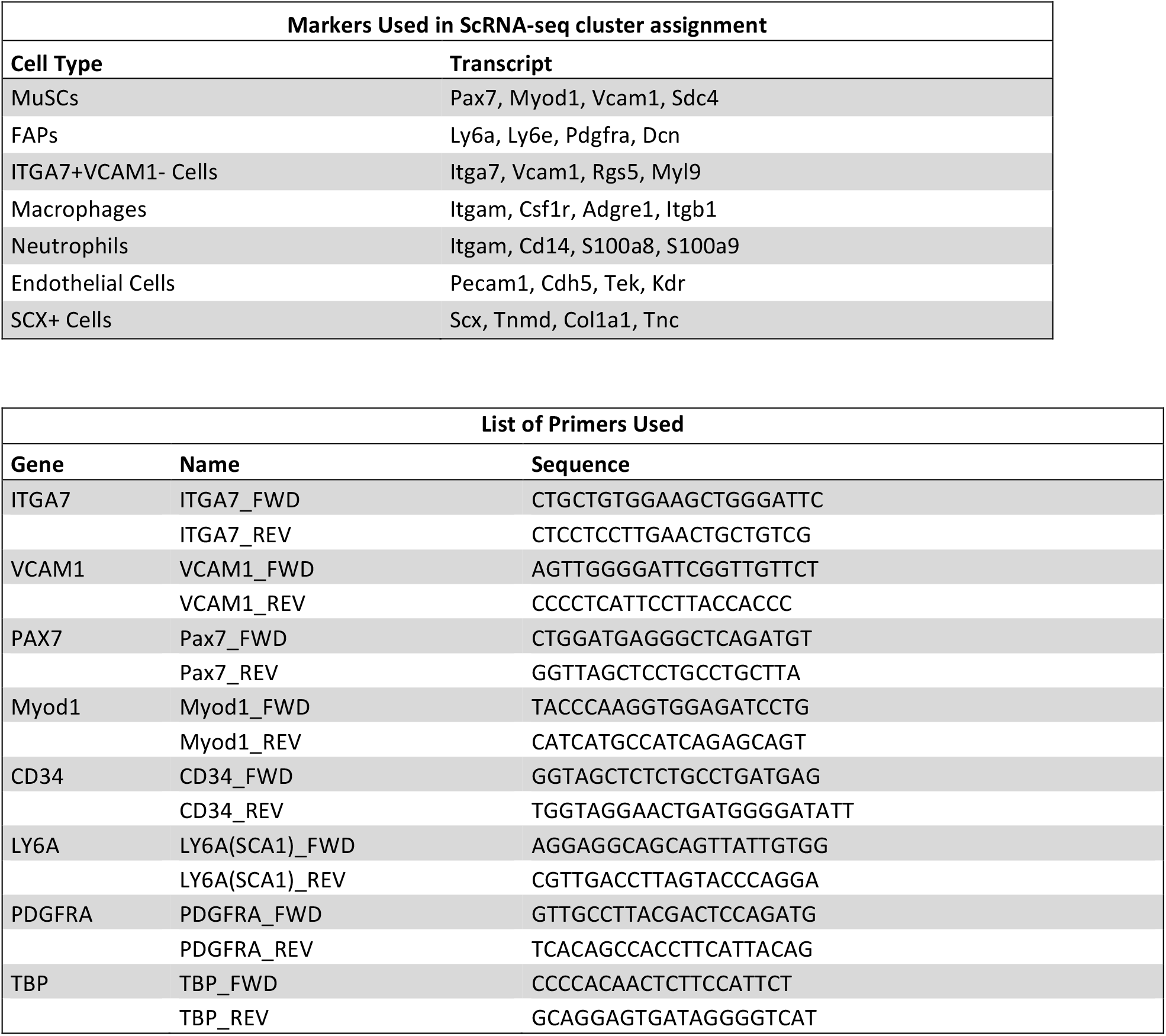
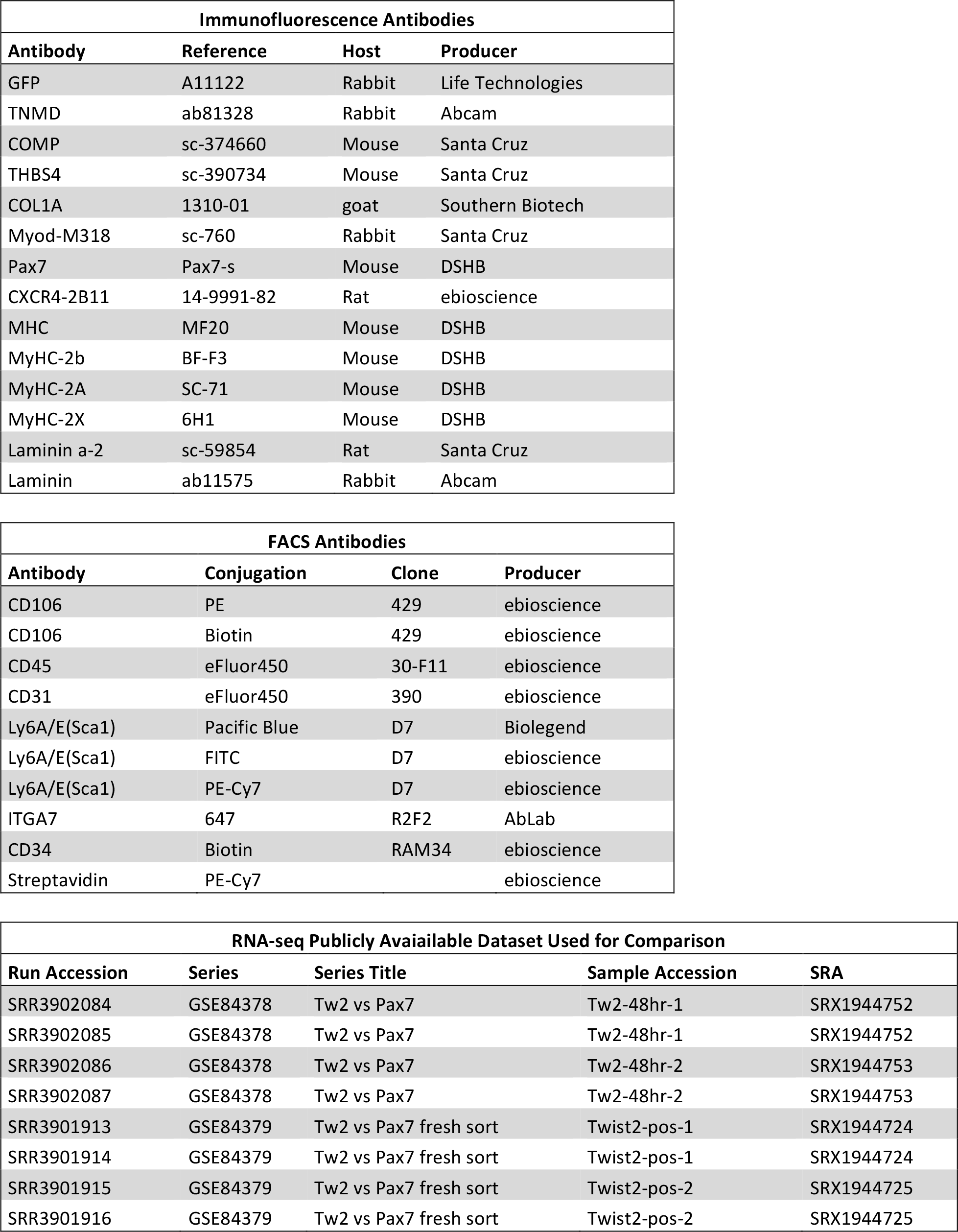
Lists of all transcripts used to assign cell population identity, primers, antibodies and publicly available datasets used in the paper.

